# Inference of multiple-wave population admixture by modeling decay of linkage disequilibrium with polynomial functions

**DOI:** 10.1101/082644

**Authors:** Ying Zhou, Kai Yuan, Yaoliang Yu, Xumin Ni, Pengtao Xie, Eric P Xing, Shuhua Xu

## Abstract

To infer the histories of population admixture, one important challenge with methods based on the admixture linkage disequilibrium (ALD) is to get rid of the effect of source LD (SLD) which is directly inherited from source populations. In previous methods, only the decay curve of weighted LD between pairs of sites whose genetic distance were larger than a certain starting distance was fitted by single or multiple exponential functions, for the inference of recent single- or multiple-wave of admixture. However, the effect of SLD has not been well defined and no tool has been developed to estimate the effect of SLD on weighted LD decay. In this study, we defined the SLD in the formularized weighted LD statistic under the two-way admixture model, and proposed polynomial spectrum (p-spectrum) to study the weighted SLD and weighted LD. We also found reference populations could be used to reduce the SLD in weighted LD statistic. We further developed a method, iMAAPs, to infer **M**ultiple-wave **A**dmixture by fitting **A**LD using **P**olynomial **s**pectrum. We evaluated the performance of iMAAPs under various admixture models in simulated data and applied iMAAPs into analysis of genome-wide single nucleotide polymorphism data from the Human Genome Diversity Project (HGDP) and the HapMap Project. We showed that iMAAPs is a considerable improvement over other current methods and further facilitates the inference of the histories of complex population admixtures.

## Introduction

The “Out of Africa” human migrations result in population differentiation in different continents, while subsequent migrations that have occurred over the past millennia led to gene flow among previously separated human sub-populations. As a consequence, admixed populations come into being when previously mutually isolated populations met and intermarry. Population admixture has received a great deal of attention recently. Many studies based on genome-wide data have shown that gene flow is common among inter-continental and intra-continental populations and that admixture of populations often leads to extended linkage disequilibrium (LD), which can greatly facilitate the mapping of human disease genes (McKeigue 2005; Reich and Patterson 2005; Smith and O’Brien 2005).

The high levels of LD are produced by admixture at loci that have different allele frequencies among the involved populations (Nei and Li 1973). Because of recombination, this particular type of admixture LD, or ALD, decays as a function of time since admixture. Consequently, it is possible to infer population admixture by modeling the dynamic changes of ALD. Moorjani et al. proposed such an approach by aggregating pairwise LD measurements through a weighting scheme (Moorjani *et al.* 2011) and its software ROLLOFF was fully explained by Patterson N. et al. (Patterson *et al.* 2012), which was further developed by Loh et al. (Loh *et al.* 2013) and by Pickrell et al (Pickrell *et al.* 2014). This ALD-based approach is particularly useful for admixture dating.

Under the hybrid isolation (HI) model, the expected value of LD decreases at the rate of 1 – *d* (Chakraborty and Weiss 1988; Pfaff *et al.* 2001), where *d* is the genetic distance (in Morgan) between two sites. And after *g* generation, the LD decays to (1 – *d*)^*g*^ of its original value, assuming that the admixed population is engaged in random mating and has infinite effective population size (Hill and Robertson 1966). Recently, Pickrell et al. considered the situation of multiple waves of admixture from different source populations and showed that LD was comprised of multiple exponential terms, each of which refers to a single admixture event (Pickrell *et al.* 2014). Zhou et.al confirmed the polynomial expression (taking *e*^−*ld*^ as approximation of (1 – *d*)^*l*^) for each wave of admixture and added the effect of source LD (SLD) from source populations into the LD’s expression under the general admixture model (Zhou *et al.* 2016). Based on this LD framework, dating admixture becomes a problem of fitting the polynomial terms in the ALD decay.

When dating admixture in empirical populations, two major factors affect the accuracy of estimation: background LD (or SLD in the context of this work) and representative reference populations. Pickrell et al presented a method based on weighted LD to deal with multiple-wave admixture. In their method, they used starting distance strategy (abandon loci whose genetic distance are shorter than a certain distance) to reduce the bias caused by SLD and they scanned global populations to find out the best pair of reference populations for each wave of admixture (Pickrell *et al.* 2014).The key assumption of their method is that the only effect by different pairs of reference populations was resulted from the relative value of exponential/polynomial coefficients of weighted LD decay. However, they neither validated this assumption nor considered the possible effect from SLD. Here, we introduced the polynomial spectrum (p-spectrum), the fitting results with polynomial functions, to reveal polynomial property of the weighted LD decay. With simulated admixed population, we confirmed the weighted LD decay curves with different pairs of source populations had similar p-spectrum and we also found that starting distance strategy could only partly reduce the SLD (Figure 3).

An alternative way to reduce SLD is to use derived source populations to estimate SLD (Zhou *et al.* 2016). Based on this idea, a new approach is developed here to infer multiple-wave admixture, which has been implemented in iMAAPs. After evaluating this method under various admixture models, we applied it to the well-known admixed populations in HGDP (Rosenberg *et al.* 2002) and HapMap (The International HapMap Consortium 2010) data and demonstrate that the current study has greatly facilitated the understanding of the admixture history of human populations.

## Materials and Methods

### Data sets

Data for simulation and empirical analysis were obtained from two public resources: Human Genome Diversity Panel (HGDP) (Rosenberg *et al.* 2002) and the International HapMap Project phase III (The International HapMap Consortium 2010). Data filtering was performed within each population with Plink (Wigginton *et al.* 2005): Samples with missing rate > 5% per individual, SNPs with missing rate > 50% and SNPs failing in Hardy-Weinberg Equilibrium test (p-value < 1 × 10^−6^) were permanently removed from subsequent analyses.

The abbreviation of populations used in this study are: YRI: The Yoruba in Ibadan, Nigeria; LWK: Luhya in Webuye, Kenya; MKK: Maasai in Kinyawa, Kenya; ASW: African Ancestry in SW USA; CEU: U.S. Utah residents with ancestry from northern and western Europe; TSI: Tuscans in Italy; MXL: Mexican Ancestry in LA, CA, USA; CHB: Han Chinese in Beijing, China; CHD: Chinese in metropolitan Denver, CO, USA. Haplotypes used as source populations in simulations are from 113 unrelated individuals of CEU and 113 unrelated individuals of YRI.

### Simulations

In order to evaluate our method in dating admixture, we employed forward-time simulations to generate haplotypes under variant admixture scenarios: HI model, two-wave model (including the cases of one donor population and two donor populations for the second wave admixture), and the model of isolation after a period of continuous admixture. Our simulations were under the framework of copying model that new haplotypes are assembled from the segments of the source populations’ haplotypes generation by generation (Li and Stephens 2003; Price *et al.* 2009), which has been used in previous work (Price *et al.* 2009). In our simulation, no mutation was considered in generating new haplotypes.

Under the HI model, the admixture events were set as having occurred 20, 50, 100, and 200 generations ago. For the two-wave (TW) model, the simulated admixed population experienced two pulses of admixture, which were at 100 and 20 generations ago, respectively, and was isolated in the remaining time. For the recent admixture in the TW model, a scenario in which only one of the source populations donated genetic materials (TW-1 model) and the other scenario where both source populations provided gene flow (TW-2 model) were simulated.

We also simulated admixed haplotypes in the scenarios of continuous migration, in which only gene flow from source populations to the admixed population is allowed and after that the admixed population is isolated to the present. In our simulation, we used modified gradual admixture (GA) (Jin *et al.* 2012) and continuous gene flow (CGF) (Pfaff *et al.* 2001) models to shape the gene flow in the migration window, which separately resulted in GA-I model and CGF-I model. Under these two models, we set the window size of migration as 80 generations and the isolation duration as 20 generations for the long last migration, and we set the window size of migration as 30 generations and the isolation duration as 70 generations for the short last migration.

Source populations also evolved in isolation so that both the reference populations and admixed population were of the same age. The sample sizes for both source populations and admixed populations were set as 5,000. More details of parameters for simulation were given in Table S1–3.

### Weighted LD statistic and its estimator under the two-way admixture model

Under the two-way admixture model (Figure 1), two source populations provide genetic materials to the newly formed admixed population. Following the notations of Zhou et. al (Zhou *et al.* 2016), the LD in the admixed population of (*n* + 1)-st generation is composed of SLD and admixture produced LD:

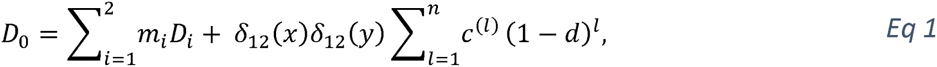
where *m*_*i*_ is the genetic proportion derived from the source population *i*, serving as the weight for linear combination of D_*i*_ (LD in source populations *i*) to form the SLD; *δ*_12_(*x*) is the allele frequency difference between population 1 and population 2 at site *x* and *d* is the genetic distance between site x and site y; *c*^(*l*)^ is a natural admixture indicator whose positive value means that admxiture happend at *l* generations ago:

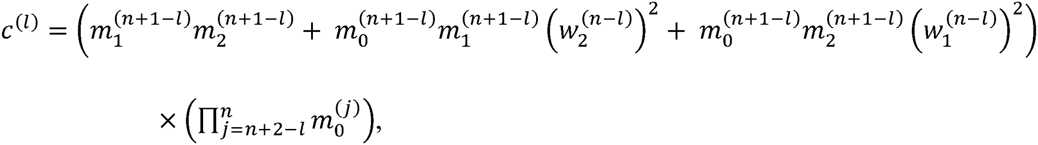
where 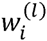 is the total genetic contribution from source population *i* in the admixed population *B*^(*l*)^.

**Figure 1.**
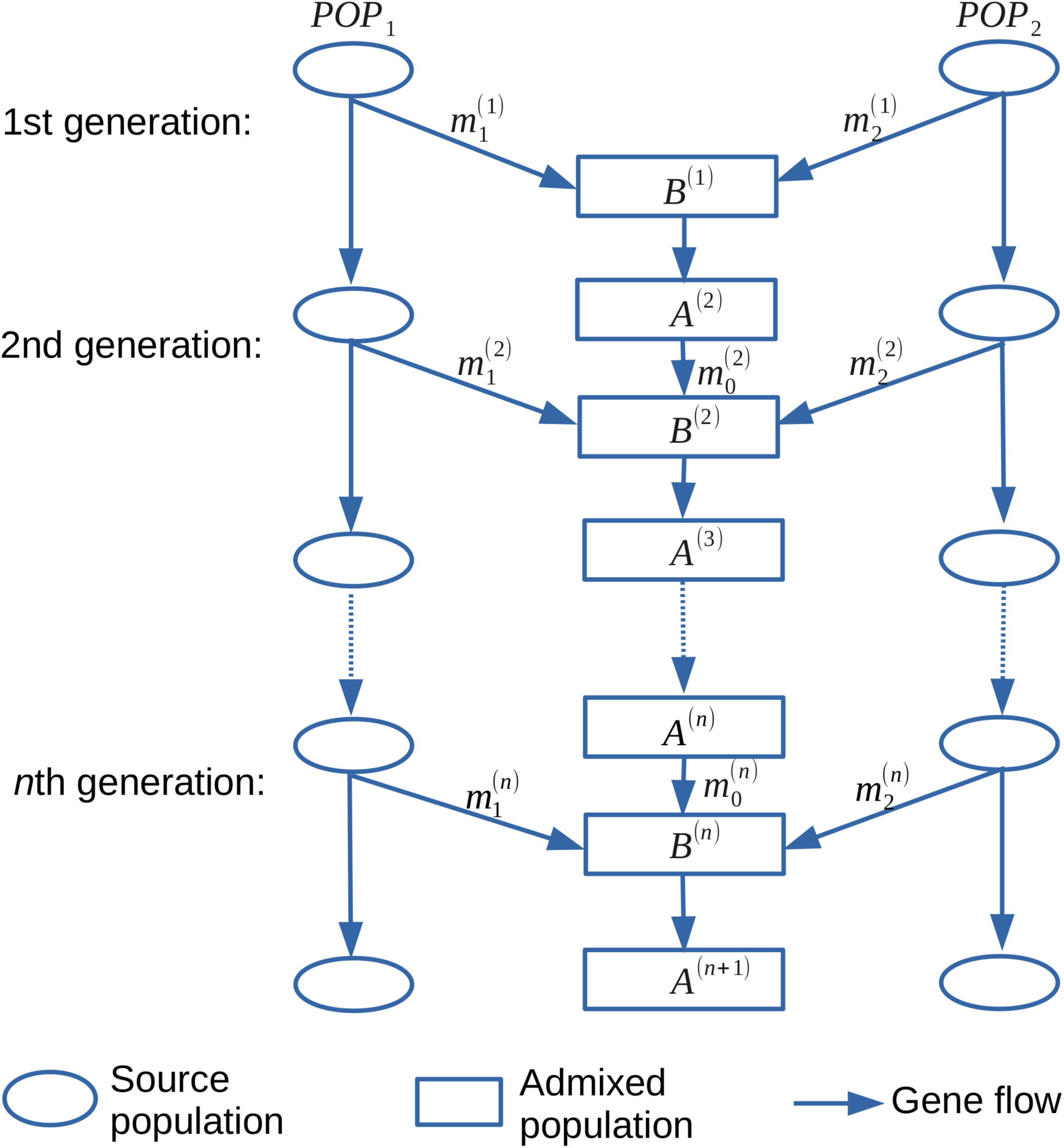
Two-way admixture model with n waves of admixture.

Using allele frequency difference *δ*_12_(*x*)*δ*_12_(*y*) as weight, the weighted LD statistic is defined as the average of LD with the weight over a set holding pairs of SNPs whose pairwise genetic distance is *d* (Loh *et al.* 2013):

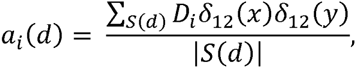
where 
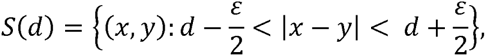
and *ε* is a discretization parameter inducing a discretization on *d.* By taking average with weight on both sides of **Error! Reference source not found.** over the set *S*(*d*), we have 
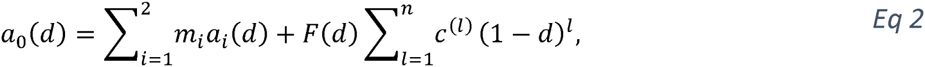
Where 
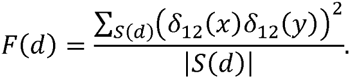

And the estimators for the weighted LD statistic for admixed population and source populations are given by Loh P. R. et. al (Loh *et al.*2013):

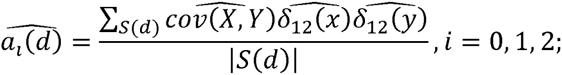

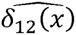 is the observed allele frequency difference and 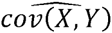 is the esimator of *D*_*i*_ between site *x* and site *y*. For the source populations, *i* = 1 or 2, and 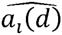 defined above is a biased estimator when the same group of samples are used for calculating both LD and the weight for the LD. Fortunately, two ways can be used for eliminating the bias: 1), divide the target population into two groups, one group is used for calculating the allele frequency difference, while the other group is used for calculating the LD (Moorjani *et al.*2011); 2), employ the unbiased statistics (Loh *et al.* 2013). In this study, we used the second method to correct the bias in the SLD estimation. Besides, *F*(*d*) can be independently estimated by:

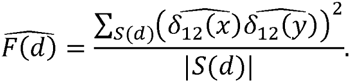

Here, we separated *F*(*d*) from the coefficients of polynomial functions to avoid the possible effect to the process of polynomial fitting.

### Factorizing of weighted LD statistic with polynomial functions

Based on formula of weighted LD statistic in the admixed population (Eq 2), admixture events are recorded in the polynomial function 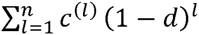, where positive value of *c*^(*l*)^ indicates the admixture at *l*th generation before present. So the direct way to date the admixture is to find out the positive value of *c*^(*l*)^. However, two possible risks may affect the results when fitting a_0_(*d*) with polynomial functions: one is *a*_*i*_(*d*), *i* = 1, 2, which represents the SLD; the other is *F*(*d*) which is a function decaying as *d* increases (Figure S3). Inspired by the Weierstrass approximation theorem, which says continuous function can be approximated by polynomial function, we used polynomial function to approximate *a*_*i*_(*d*), *i* = 0, 1, 2, and *F*(*d*) to explore the posssible interaction between them. Actually, by fitting the decay curve with polynomial function 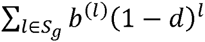, we got the spectrum that the *b*^(*l*)^ values on set *S*_*g*_, which we defined as *polynomial spectrum* (p-spectrum) (Figure 3). In the polynomial fitting, *b*^(*l*)^ needs to be non-negative and *S*_*g*_ is a finite set holding the candidate time points for the possible admixture signals. This numeric method to generate the p-spectrum is illustrated in ***Appendix**.* Replacing *a*_*i*_(*d*),*i* = 1,2 and *F*(*d*) with polynomial functions 
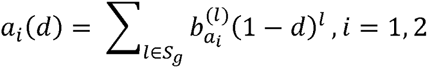

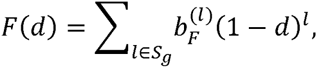
*a*_0_(*d*) turns to be 
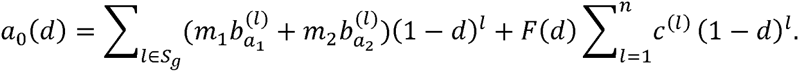

This expression of *a*_0_(*d*) tells us that SLD, linear combination of *a*_*i*_(*d*), *i* = 1, 2, would bring in false positive admixture signals while *F*(*d*) has the potential to destroy the admixture time inference when we try to fit *a*_0_(*d*) directly with polynomial functions. So it is essential to evaluate the effect from *a*_*i*_(*d*), *i* = 1, 2, and *F*(*d*). Fortunately, *a*_*i*_(*d*), *i* = 1, 2, and *F*(*d*) can be estimated with the source populations so that the effect can be evaluated.

To evaluate the effect from *a*_*i*_(*d*), *i* = 1, 2, and *F*(*d*), we simulated a 100-generation-old admixed population under HI model and the simulated admixed population was initiated with the haplotype of YRI and CEU of the proportion 50%:50%. Derived source populations of YRI and CEU were also generated, separately. Based on the simulated genotype data in both source populations and admixed population, both 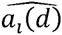 and 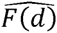 can be calculated through a Fast Fourier Transform algorithm, which is able to increase the computational efficiency (Loh *et al.* 2013). After that, the p-spectrum was constructed on a time set ranging from 0 to 2000 generations accordingly. In the spectrum of 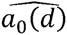, we found three bunches of signals: two sharp bunches appeared around 100 and 1,250 generations and one flat bulb lay around 180 generations (Figure 2). Both in the spectrum of 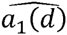 and the spectrum of 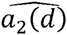, we found signals around the value 1,250 generations and signals close to 250 generations (Figures S1, S2). And in the p-spectrum of 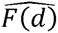, we found only a strong peak on the time 0 and two weak signal peaks over 250 generations ago, which explained the sharp decay in its decay curve (Figures S3), suggesting that we need to consider this effect to precisely resolve admixture. In the time spectrum of 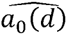, signals around 100 could be explained easily by the designed admixture and both signals around 1250 and signals around 180 are probably introduced by the SLD. To test this explanation, we directly constructed *z*(*d*) as

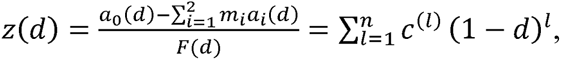

which can be estimated with the simulated admixed population and derived source populations by 
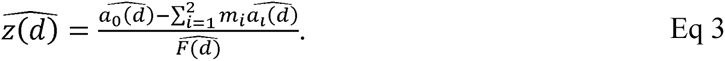

In the p-spectrum of 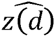, the relative strength of noise-like signals outside the bunch of signals around 100 generation become much weaker than that in the p-spectrum of 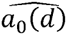 (Figure S4). This result confirms our explanation on the p-spectrum of 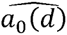 and indicates source populations can be used to reduce effect of SLD.

**Figure 2.**
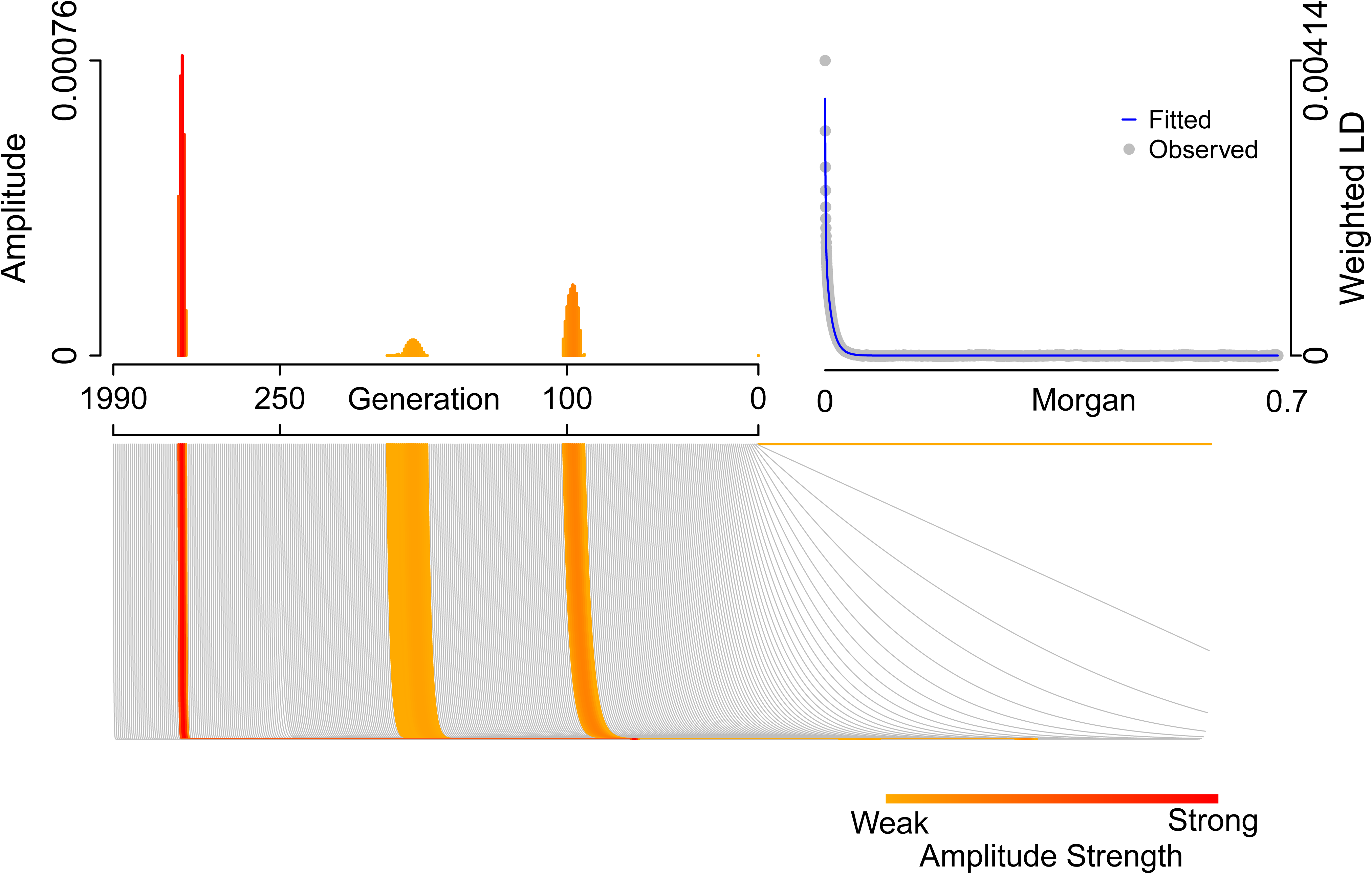
p-spectrum for 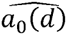 in a simulated admixed population. The observed weighted LD decay (gray points at top right) are fitted by hundreds of polynomial functions (gray curves in the bottom panel), and a few of them whose coefficients are positive (highlighted in heat color) and the amplitude for each positive coefficient are plotted along the value of 1 (generation before present) at top left.

### Time inference for multiple-wave admixture

Because *c*^(*l*)^ is a natural indicater for admixture event, the natural extension for the p-spectrum is to infer the admixture time. In this section, we are going to present the time inference method based on p-spectrum. In empirical populations, both the 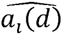, *i* = 0, 1, 2 and 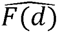 can be calculated based on the genotype data of the admixed population and reference populations. Meanwhile, the population admixture proportions are estimated by 
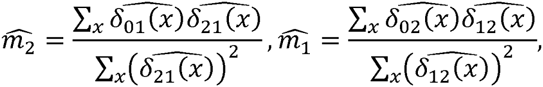
then 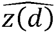 can be calculated so as its p-spectrum 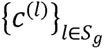.

Next, we are going to date admixture and evaluate the existence for each wave of admixture with a Jackknife-based method. Suppose we have 22 autosomes for the target admixed population, and each chromosome is excluded one at each time to calculate decay curve of 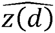, (Loh *et al.* 2013). This means when chromosome *i* is excluded, the remaining 21 chromosomes are used to calculate 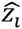 and the p-spectrum 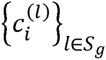. Then the p-values are attained on each *l* by testing whether the values in the set 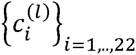 is bigger than zero, and we used 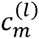, the median of 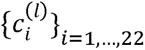,as the summary p-spectrum for the target population. We could also use the mean value to construct 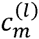, but it would lead to more false admixture signals. Based on the summary p-spectrum, *l* with positive values of 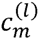 are gathered as the candidate admixture time points and then they are clustered into groups as different waves of admixture, say 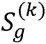 for kth wave of admixture. Once these time points are grouped, the mean and variance of the time for that wave of admixture can be calculated by 
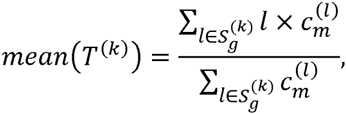

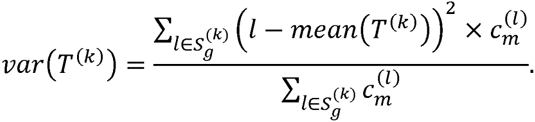

Meanwhile, we used the minimum p-value on each time point in that group to measure the significance for each wave of admixture. In this way, we could date multiple-wave admixture and measure the significance of each wave of admixture. This method has been implemented in the Software named iMAAPs, which is available at http://www.picb.ac.cn/PGG/resource.php

## Results

### Dating multiple-wave admixture with weighted LD statistic

There are two main difficulties for dating admixture in empirical analysis: reference populations selection and SLD reduction. To deal with these two difficulties, Pickrell et. al claimed that different pairs of reference populations often have different relative values but always have the same sign of the coefficient of (1 − *d*)^*l*^ so that they can traverse all pairs of reference populations to test the presence for the possible admixture and estimate the time for each wave of admixture. And they used the LD whose pairwise genetic distance is longer than 0.5 centiMorgans (cM), which was supposed to reduce the effect of SLD. Meanwhile, they also claimed that their algorithm was not very powerful in detecting multiple admixture (Loh *et al.* 2013; Pickrell *et al.* 2014). Under our framework of weighted LD (Eq 2), we confirmed that *c*^(*l*)^ is an admixture determined constant that it is independent to the selection of reference populations; we pointed out both the *a*_*i*_(*d*), *i* = 1, 2 and *F*(*d*) have the potential to affect the p-spectrum of *a*_0_(*d*), which directly affect the estimation of the coefficient of (1 − *d*)^*l*^; we also noticed that the effect of SLD reduction with starting distance was also not evaluated in Pickrell et. al’s work, which may be the reason that their method is not so powerful in detecting multiple admixture. To verify our conjecture, we constructed the summary p-spectrum for weighted LD decay curves on a simulated admixed population with different pairs of reference populations.

A 100-generation-old admixed population was generated under HI model, with YRI and CEU as source populations of admixture proportion 0.5:0.5. Total 55 pairs of HapMap populations (YRI, LWK, MKK, ASW, CEU, TSI, MXL, CHB, CHD, GIH, JPT) were used as references to calculate weighted LD *a*_0_(*d*) for further p-spectrum construction. In the summary, p-spectrum with full weighted LD decay (Figure 3A) for nearly all pairs of reference populations arose three main peaks around 100, 180 and 1250 generations. In the p-spectrum with weighted LD decay began at 0.5 cM (Figure 3B), the peak around 180 and 1250 generations disappeared but a new peak around 120 generations arose, which was probably the remaining SLD and it may bias the time estimation of admixture. The remaining SLD should be the reason why ALDER did not work well for multiple admixture. Meanwhile, we also observed that weighted LD decay with the pairs of reference populations close to the true source populations would have similar p-spectrum to what we want to reveal (Figure 3), which indicated us to use populations not the exact but similar to the source populations as references to construct the p-spectrum. This observation also supported that using proper reference populations could increase the accuracy of ALDER in resolving weighted LD decay.

**Figure 3.**
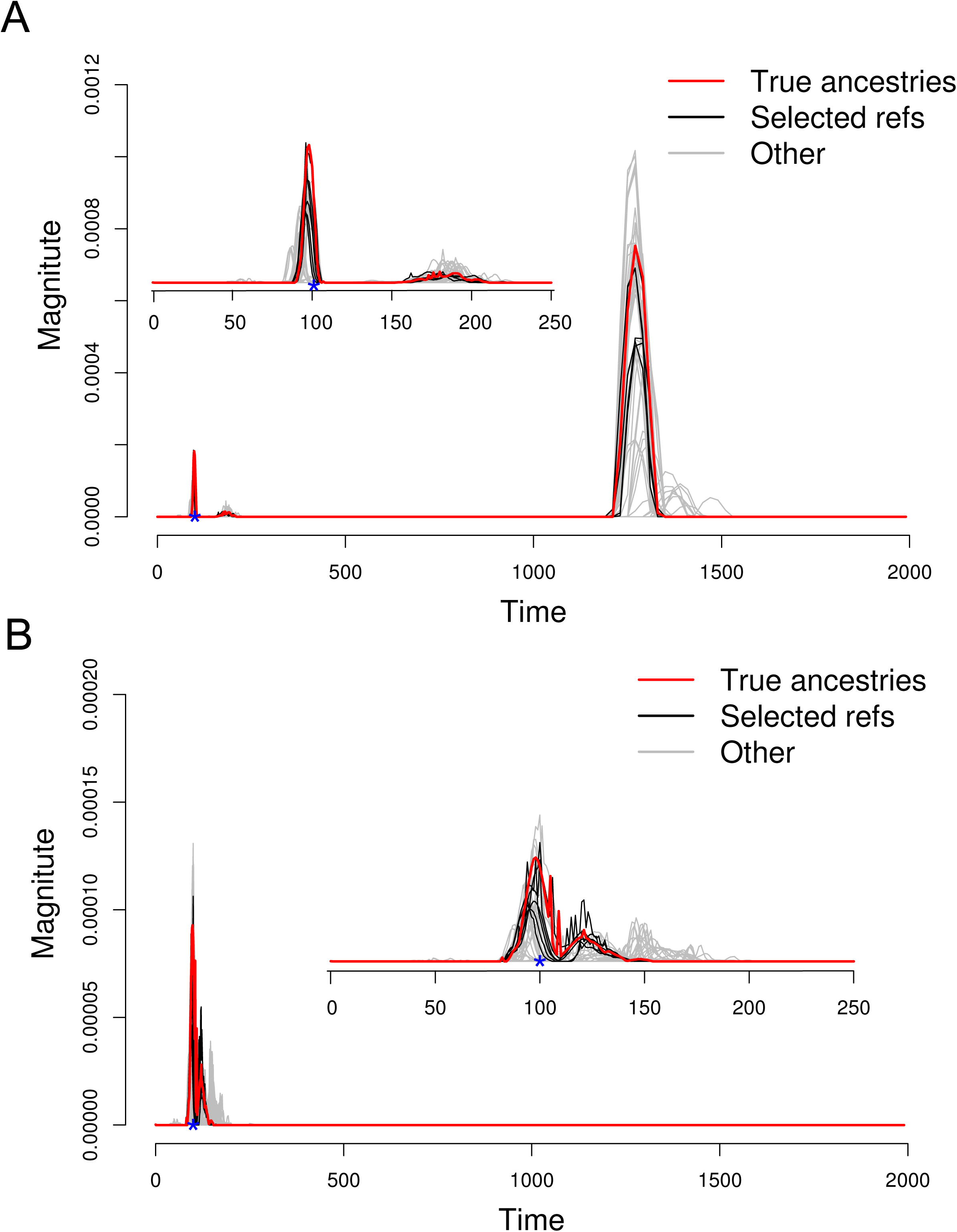
Summary p-spectrum for 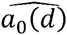 with all pairs of populations from HapMap. Spectrum with true source populations (CEU, YRI) is in red lines; spectrum with selected pairs of reference populations (CEU, LWK; CEU, MKK; TSI, LWK; TSI, YRI; TSI, MKK;) are in black lines; spectrums with other pairs of reference populations are in gray lines. **A**: Summary p-spectrum for full LD decay; **B** Summary spectrum for LD decay of starting distance 0.5 cM.

### Evaluation of iMAAPs

In our p-spectrum based method iMAAPs, we used reference populations to estimate SLD and *F*(*d*), and separated their effect from the weighted LD of admixed population. Thus, we could directly estimate the admixture determined parameters *c*^(*l*)^ and the time and the waves of admixture. A workable method in empirical admixture analysis should be robust to the proxy source populations. We have observed the robustness of p-spectrum to different pairs of reference populations, so we will evaluate the performance of iMAAPs to different reference pairs. Here, with the simulated 100-generation-old African-European admixed population, generated by YRI and CEU, we showed that iMAAPs is very robust with African-European pairs (YRI, CEU; LWK, CEU; MKK, CEU; LWK, TSI; YRI, TSI; MKK, TSI) as reference populations to infer the admixture time (Table S4).

We also tested our method under various admixture model, iMAAPs is able to reconstruct the history of the admixture population well. For the one-pulse and two-pulse admixture models, iMAAPs gave the time close to the true admixture; for the continuous migration models, it was able to place most of the signals in a particular migration time window (Figure 4, Figure S5–13).

**Figure 4.**
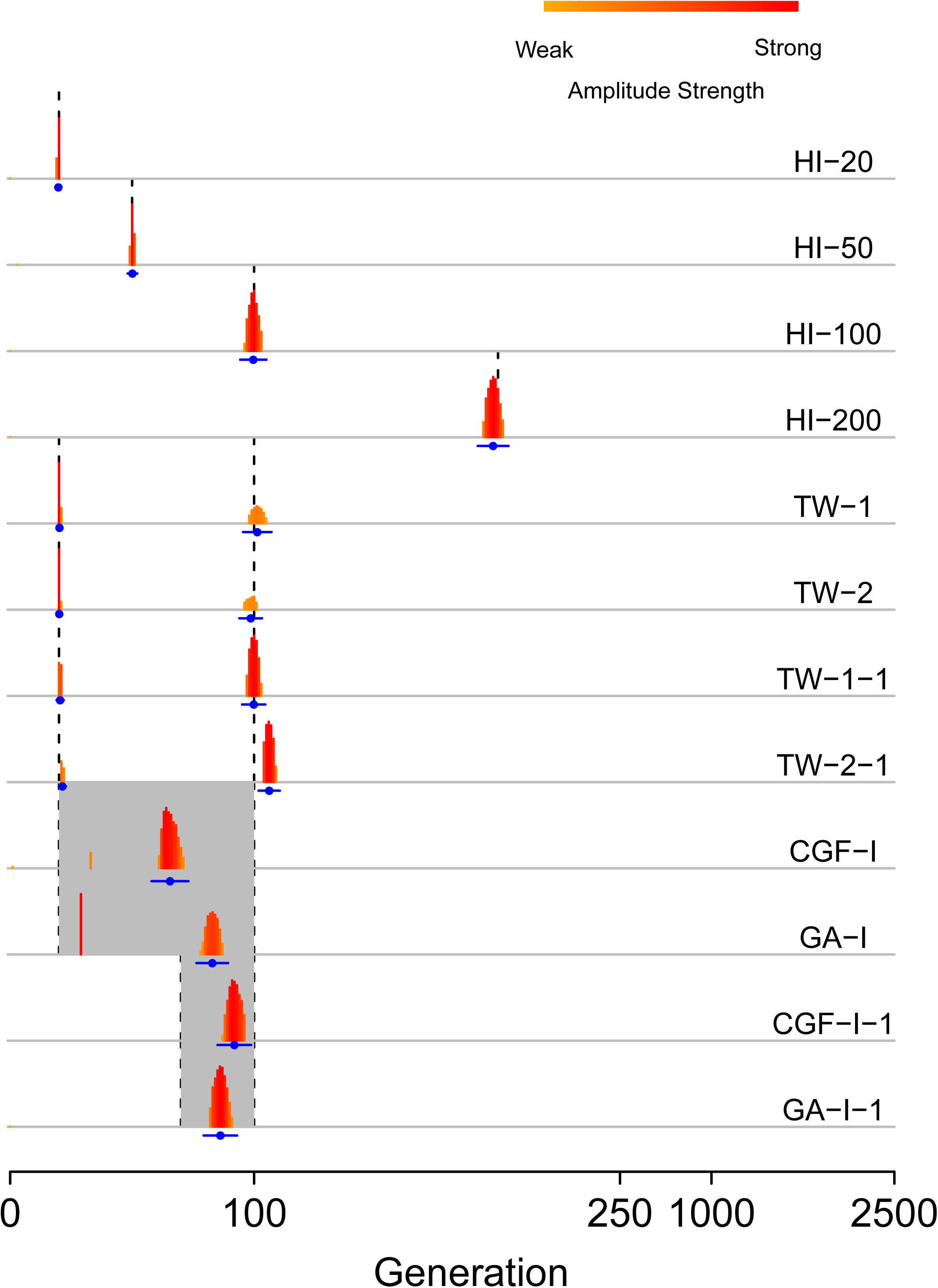
The performance of iMAAPs under various admixture models. The black vertical dash lines represent the true simulated admixture time and gray areas represent the time window for continuous admixture. The summary p-spectrum of 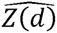 for each simulated admixed population is plotted in heat color and the estimated admixture times are plotted in blue points, the mean values, and lines, the ranges of 3 times of standard derivation.

### Empirical analysis

The current method was first applied to a few well-known admixed populations from available public databases: HGDP (Rosenberg *et al.* 2002) and HapMap Project phase III (The International HapMap Consortium 2010). Our method is currently designed under the framework of two-way admixture and source populations or the populations similar to which are required in empirical analysis. Besides, two principles should be considered for interpretation:

(1) Existence of estimations for longer than 500 generations indicates the SLD has not been well removed so that some of the admixture signals, especially for ancient ones, are probably generated by the SLD instead of the admixture.

(2) Existence of estimations close to generation 1 is always considered as the result of the population substructure but not the admixture.

Based on these principles, we first analyzed three well-known admixed populations: African American (57 ASW individuals from HapMap), Mexican (86 MXL individuals from HapMap) and Uygur (10 Uygur individuals from HGDP). We also used ALDER to analyze these admixed populations (Table S5). For each admixed population, we conducted three rounds of estimations. In the first round, we used all the populations in the full data set as the references to infer the admixture; in the second round, we used population pairs with the highest amplitude for each wave of admixture in the first round as the reference populations to re-run ALDER; in the last round, we selected populations according to the admixture pattern based on the population inference in the first round. That is to say, if CEU and YRI are inferred as the best pair of populations to explain the admixture, then we selected all populations that could represent European ancestries and African ancestries as reference populations in the third run of ALDER, which we believed would increase the estimation accuracy.

In our analysis with iMAAPs, reference populations were selected based on the results of ALDER’s inference on each wave of admixture. CEU (n = 113) and YRI (n = 147) were chosen as the ancestral populations of ASW. YRI (n = 147), TSI (n = 102) and American Indian (7 Colombians, 14 Karitiana, 21 Maya, 14 Pimas and 8 Suruis from HGDP) were used as the ancestral populations of MXL. Basque (n = 24), Sardinian (n = 28), Japanese (n = 28), Han (n = 34) and French (n = 28) were used as the ancestral populations of Uygur.

The estimation of admixture time for ASW is 5.4+/-0.4 generations ago and the SLD has been well reduced with YRI and CEU as reference populations (Table 1, Figure S15). In the meantime, ALDER gives us two different results: 12.0+/-4.4 generations with all populations in HapMap as references; 6.3+/-3.3 and 77.0+/-65.9 generations with selected reference populations from HapMap (Table S4). In this estimation of ALDER, generation 6.3 is very close to our result, which can be interpreted as the admixture time of the population ASW. And the result on generation 77.0 reflects the failure of SLD reduction with starting distance of 0.5 cM.

**Table 1.** Time of admixture (generations) determined using iMAAPs with selected reference populations. In each time cell, the mean value ± standard derivation is listed with P-value for the significance of existence of that wave of admixture.

| Admixed population | Reference populations | Time 1 | Time 2 | Time 3 | Time 4 |
| --- | --- | --- | --- | --- | --- |
| **ASW** | CEU;YRI | 0.0±0.0 | 5.4±0.4 |  |  |
| | | $3.92 \times 10^{-40}$ | $4.15 \times 10^{-17}$ | | |
| **MXL** | American Indian;TSI | 0.1±0.3 | 7.0±0.2 |  |  |
| | | $8.69 \times 10^{-27}$ | $1.70 \times 10^{-13}$ | | |
| **MXL** | American Indian;YRI | 0.0±0.0 | 8.2±0.4 |  |  |
| | | $1.82 \times 10^{-31}$ | $8.15 \times 10^{-19}$ | | |
| **MKK** | CEU;YRI | 1.0±0.0 | 16.2±0.38 | 68.4±1.5 | 653.0±15.0 |
| | | $1.33 \times 10^{-28}$ | $1.10 \times 10^{-4}$ | $5.47 \times 10^{-5}$ | $3.43 \times 10^{-20}$ |
| **MKK** | TSI;YRI | 1.1±0.3 | 17.9±0.23 | 70.3±1.4 | 632.1±15.1 |
| | | $5.26 \times 10^{-28}$ | $7.11 \times 10^{-5}$ | $4.29 \times 10^{-5}$ | $3.36 \times 10^{-13}$ |
| **Uygur** | Basque;Han | 0.0±0.0 | 2.4±0.5 | 30.0±0.0 |  |
| | | $4.37 \times 10^{-4}$ | $4.22 \times 10^{-7}$ | $2.48 \times 10^{-6}$ | |
| **Uygur** | French;Han | 0.0±0.0 | 4.4±0.5 | 33.3±0.5 |  |
| | | $5.81 \times 10^{-12}$ | $4.54 \times 10^{-5}$ | $6.23 \times 10^{-5}$ | |
| **Uygur** | Sardinian; Japanese | 2.8±0.1 | 34.0±0 |  |  |
| | | $9.71 \times 10^{-12}$ | $7.65 \times 10^{-4}$ | | |

MXL seems to have experienced the main admixture of 7.0+/-0.2 generations ago with TSI and American Indian as reference populations; 8.2+/-0.4 generations ago with YRI and American Indian as reference populations (Table 1). More admixture time points would be detected using the mean to construct summary p-spectrum (Table S5), which needs to be confirmed by further studies.

The Uygur population has been reported to have much longer admixture history than ASW and MXL (Xu and Jin 2008; Xu *et al.* 2008; Qin *et al.* 2015). The admixture is resolved at 33.3+/-0.5 generations ago with Han and French as reference populations. This admixture event has also been detected with Basque, Han, Sardinian and Japanese as reference populations, suggesting that the major admixture formatting ALD in the present population happened around 825 years ago.

Loh et al. speculated that there could have been multiple waves of admixture in the history of MKK (Loh *et al.* 2013). Here, both our method and ALDER detected at least two waves of admixture (Table 1, Table S4, and Figure S15). We used reference pair of YRI and CEU and pair of YRI and TSI to resolve the admixture of the MKK. With YRI and CEU as references, admixtures around 16.2 and 68.4 generations ago were detected; with YRI and TSI as references, admixtures around 17.9 and 70.3 generations ago were detected. However, both detections have estimations longer than 500 generations ago, indicating that we need to take care of SLD when interpreting the time of admixture.

## Discussion

Available methods based ALD for admixture dating have shown their robustness in dealing with genotype data (Loh *et al.* 2013) and complicated admixture history (Pickrell *et al.* 2014). However, the effect of SLD in these methods has not been well defined and efficiently reduced, which may bias the estimation. In this study, we defined the SLD in the weighted LD statistic of the target admixed population under two-way admixture model, and introduced the p-spectrum to study the weighted LD decay for both source populations and admixed population. We found that SLD pretends to have higher degree of p-spectrum than the LD introduced by recent admixture and using starting distance can partly reduce the effect of SLD (Figure 3). We also found that SLD can be well compensated by the LD of source populations (Figure S5–S13), which motivated us to develop a new method iMAAPs to infer multiple wave admixture. In this method, we used reference populations to estimate the SLD and then reduced its effect and gave the accurate estimation.

We applied both iMAAPs and ALDER to date several well-known admixed populations. When running ALDER, we conducted estimation in three rounds with different sets of reference populations. Based on the ALDER’s assumption of the effect of different pairs of reference populations on weighted LD, time estimations in these three rounds should be much close to each other. However, for the population of ASW, the result of first round is different from the results of other rounds, indicating the potential risk for using ALDER with global references. In the estimations of the second and third round, we found signals around 6 and 70 generations before present, and the signal around 70 generations should be the false admixture signal caused by the remaining SLD, because iMAAPs only detected the signal close to the 6 and the SLD was well reduced. And we found signals were close to 0 in all these three rounds of estimation, which, as we suggested, should be interpreted as population substructure instead of admixture time. We also ran iMAAPs in populations of MXL, Uygur, and MKK. We found MXL seems to experience admixture at 7 generations ago. For the Uygur, the major admixture happened at about 33 generations ago.This date is a little ealier than the date with ALDER with three rounds, in which the most significant admixture is around 40 generations ago. The difference is probably caused by the remaining SLD with starting distance strategy. Both ALDER and iMAAPs confirmed the estimation that the population MKK experienced multiple waves of admixture, which had been predicted in the work of Loh et al. (Loh *et al.* 2013)

One of the fundamental idea in this algorithm is to take advantage of proper representative reference populations to reduce the effect of SLD. Since using improper reference populations may bias the final estimation, it is crucial to carefully select reference populations in empirical analysis. Another issue that should be noticed is that this algorithm might give mulitple pulses of signals even though the true population history was long-last conituous admixture (Figure S5–8). Nevertheless, this work built the framework of weighted LD under two-way admixture and provided an alternative way to reduce the effect of SLD in estimation of admixture time.

## Author contributions

Conceived and designed the study: **SX**. Led this study: **YZ**. Developed methods and computer tools: **YZ KY**. Analyzed the data: **YZ** and **KY**. Interpreted the data and wrote the paper: **SX YZ KY YY XN PX EX**.

## Acknowledgements

We are grateful to Dr. Li Jin, Dr. Yungang He, Dr. XiaoMing Liu and Dr. Minxian Wang for their valuable suggestions. This work is supported by the Strategic Priority Research Program (XDB13040100) and Key Research Program of Frontier Sciences (QYZDJ-SSW-SYS009) of the Chinese Academy of Sciences (CAS), the National Natural Science Foundation of China (NSFC) grant (91331204), the National Science Fund for Distinguished Young Scholars (31525014), the Program of Shanghai Academic Research Leader (16XD1404700), and the National Key Research and Development Program (2016YFC0906403). S.X. is Max-Planck Independent Research Group Leader and member of CAS Youth Innovation Promotion Association. S.X. also gratefully acknowledges the support of the National Program for Top-notch Young Innovative Talents of The "*Wanren Jihua*" Project. The funders had no role in study design, data collection and analysis, decision to publish, or preparation of the manuscript.

## Appendix Construct the p-spectrum for decay curve

The decay curve of *z*(*d*) can be fitted by a numerical routine known as the proximal gradient (Beck A. and Teboulle 2009) to minimize the objective function to fit out the parameter *c*: 
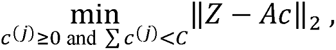
where *Z* = (*z*(*d*_1_)), *z*(*d*_2_), …, *z*(*d*_*s*−1_), *z*(*d*_*s*_))^*T*^is a vector of *z*(*d*) with different *d* values. *c* = (*c*^(0)^, *c*^(1)^, …, *c*^(*n*−1)^, *c*^(*n*)^)^*T*^ is the coefficient of the polynomial functions and all entries in vector *c* are non-negative. The (*i*, *j*)*th* entry of the matrix *A*_*s*×(*n*+1)_ is *A*_*ij*_ = (1 − *d*_*i*_)^*j*^, where *j* = 0,1,..,*n*. In empirical analysis, the value of *j* ranges from 0 to thousands, say 0 to 2000 in our analysis, which would lead the matrix *A*_*ij*_ to be too large to computate effeciently. In order to increase the computation efficiency, we use the set *S*_*g*_, a subset containing time candidates sampled from the range of 0 to 2000, as the *j*’s value set. With this method, it was possible to find the vector *c* so as the fitting curve *Ac*. Next, we construct the p-spectrum from denoising the vector *c*.

For the vector *c*, each entry represents the magnitude of the signal and large magnitude indicates that the signal contains information rather than noise. Based on this idea, only the top *c*^(*l*)^ values that together composed 99.9% of *Z* were retained for the p-spectrum construction. This means that we need to find out the smallest set *Ω* ⊆ *S*_*g*_ subject to the condition 
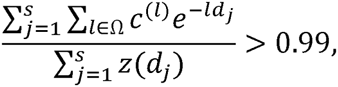
and then let *c*^(*l*)^, *l* ∈ *S*_*g*_\Ω to be zero. In this way, we constructed the p-spectrum for the decay curve *Z*.

### Clustering the candidate time points

Suppose we have the candidate time points generated from summary p-spectrum as increasing series {*t*_*i*_}_*i*_=l,2,…,*I* then we say *t*_*i*_ and *t*_*i*+1_ belong to the same cluster if only one of these two conditions stands: 
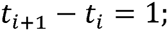
or 
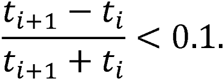

**Figure S1:** p-spectrum for 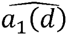(d) in one of the reference populations (CEU). The observed weighted LD decay (gray points at top right) are fitted by hundreds of polynomial functions (gray curves in the bottom panel), and a few of them whose coefficients are positive (highlighted in heat color) and the amplitude for each positive coefficient are plotted along the value of *l* (generation before present) at top left.

**Figure S2:** p-spectrum for 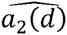 in one of the reference populations (YRI). The observed weighted LD decay (gray points at top right) are fitted by hundreds of polynomial functions (gray curves in the bottom panel), and a few of them whose coefficients are positive (highlighted in heat color) and the amplitude for each positive coefficient are plotted along the value of *l* (generation before present) at top left.

**Figure S3:** p-spectrum for 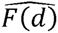. The observed 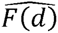 values (gray points at top right) are fitted by hundreds of polynomial functions (gray curves in the bottom panel), and a few of them whose coefficients are positive (highlighted in heat color) and the amplitude for each positive coefficient are plotted along the value of *l* (generation before present) at top left.

**Figure S4:** p-spectrum for 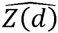. The observed 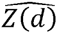 values (gray points at top right) are fitted by hundreds of polynomial functions (gray curves in the bottom panel), and a few of them whose coefficients are positive (highlighted in heat color) and the amplitude for each positive coefficient are plotted along the value of *l* (generation before present) at top left.

**Figure S5:** The performance of iMAAPs on 10 independent simulated admixed populations under CGF-I model (case 1). The gray area represents the time window for continuous admixture. The summary p-spectrum of 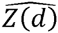 for each simulated admixed population is plotted in heat color. The blue points represent the mean values, and the blue lines represent the ranges of 3 times of standard derivation.

**Figure S6:** The performance of iMAAPs on 10 independent simulated admixed populations under CGF-I model (case 2). The gray area represents the time window for continuous admixture. The summary p-spectrum of 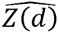 for each simulated admixed population is plotted in heat color. The blue points represent the mean values, and the blue lines represent the ranges of 3 times of standard derivation.

**Figure S7:** The performance of iMAAPs on 10 independent simulated admixed populations under GA-I model (case 1). The gray area represents the time window for continuous admixture. The summary p-spectrum of 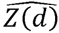 for each simulated admixed population is plotted in heat color. The blue points represent the mean values, and the blue lines represent the ranges of 3 times of standard derivation.

**Figure S8:** The performance of iMAAPs on 10 independent simulated admixed populations under GA-I model (case 2). The gray area represents the time window for continuous admixture. The summary p-spectrum of 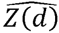 for each simulated admixed population is plotted in heat color. The blue points represent the mean values, and the blue lines represent the ranges of 3 times of standard derivation.

**Figure S9:** The performance of iMAAPs on 10 independent simulated admixed populations under HI model. The black vertical dash lines represent the true simulated admixture time. The summary p-spectrum of 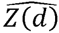 for each simulated admixed population is plotted in heat color. The blue points represent the mean values, and the blue lines represent the ranges of 3 times of standard derivation.

**Figure S10:** The performance of iMAAPs on 10 independent simulated admixed populations under TW-1 model (case 1). The black vertical dash lines represent the true simulated admixture time. The summary p-spectrum of 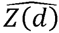 for each simulated admixed population is plotted in heat color. The blue points represent the mean values, and the blue lines represent the ranges of 3 times of standard derivation.

**Figure S11:** The performance of iMAAPs on 10 independent simulated admixed populations under TW-1 model (case 2). The black vertical dash lines represent the true simulated admixture time. The summary p-spectrum of 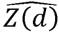 for each simulated admixed population is plotted in heat color. The blue points represent the mean values, and the blue lines represent the ranges of 3 times of standard derivation.

**Figure S12:** The performance of iMAAPs on 10 independent simulated admixed populations under TW-2 model (case 1). The black vertical dash lines represent the true simulated admixture time. The summary p-spectrum of 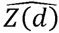 for each simulated admixed population is plotted in heat color. The blue points represent the mean values, and the blue lines represent the ranges of 3 times of standard derivation.

**Figure S13:** The performance of iMAAPs on 10 independent simulated admixed populations under TW-2 model (case 2). The black vertical dash lines represent the true simulated admixture time. The summary p-spectrum of 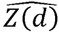 for each simulated admixed population is plotted in heat color. The blue points represent the mean values, and the blue lines represent the ranges of 3 times of standard derivation.

**Figure S14:** The performance of iMAAPs on empirical admixed populations. The summary p-spectrum of 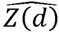 for each simulated admixed population, calculated with the median, is plotted in heat color. The blue points represent the mean values, and the blue lines represent the ranges of 3 times of standard derivation.

